# Toll-like receptor 4 is activated by platinum and contributes to cisplatin-induced ototoxicity

**DOI:** 10.1101/2020.06.19.162057

**Authors:** Ghazal Babolmorad, Asna Latif, Niall M. Pollock, Ivan K. Domingo, Cole Delyea, Aja M. Rieger, W. Ted Allison, Amit P. Bhavsar

## Abstract

Toll-like receptor 4 (TLR4) is famous for recognizing the bacterial endotoxin lipopolysaccharide (LPS) as its canonical ligand. TLR4 is also activated by other classes of agonist including some Group 9/10 transition metals. Roles for these non-canonical ligands in pathobiology mostly remain obscure, though TLR4 interactions with metals can mediate immune hypersensitivity reactions. In this work, we tested whether TLR4 can be activated by the Group 10 transition metal, platinum. We demonstrated that in the presence of TLR4, platinum activates pathways downstream of TLR4 to a similar extent as the known TLR4 agonists LPS and nickel. Platinum is the active moiety in cisplatin, a very potent and invaluable chemotherapeutic used to treat solid tumors in childhood cancer patients. Unfortunately, cisplatin use is limited due to an adverse effect of permanent hearing loss (cisplatin-induced ototoxicity, CIO). Herein, we demonstrated that cisplatin also activates TLR4, prompting the hypothesis that TLR4 mediates aspects of CIO. Cisplatin activation of TLR4 was independent of the TLR4 co-receptors CD14 and MD-2, which is consistent with TLR4 signaling elicited by transition metals. We found that TLR4 is required for cisplatin-induced inflammatory, oxidative and apoptotic responses in an ear outer hair cell line and for hair cell damage *in vivo*. Thus, TLR4 is a promising therapeutic target to mitigate CIO. We additionally identify a TLR4 small molecule inhibitor able to curtail cisplatin toxicity *in vitro*. Further work is warranted towards inhibiting TLR4 as a route to mitigating this adverse outcome of childhood cancer treatment.

**Significance Statement:** This work identifies platinum, and its derivative cisplatin, as new agonists for TLR4. TLR4 contributes to cisplatin-induced hair cell death *in vitro* and *in vivo*. Genetic and small molecule inhibition of TLR4 identify this receptor as a druggable therapeutic target with promise to curtail cisplatin-induced ototoxicity, a devastating side-effect of an otherwise invaluable chemotherapeutic tool.

## Introduction

Toll-like Receptor 4 (TLR4) is a membrane-bound pattern recognition receptor that is best characterized for its ability to initiate innate immune signaling upon detection of the gram-negative bacterial surface component lipopolysaccharide (LPS) (1). LPS detection requires the TLR4 co-receptors CD14 and MD-2. Structural analyses have revealed that the LPS binding pocket is comprised of both MD-2 and TLR4 on the external face of the membrane, with MD-2 making a major contribution to agonist binding (2, 3). LPS binding to the TLR4/MD-2 complex induces TLR4 dimerization and signal propagation through adapter protein recruitment on the cytoplasmic face of the membrane (2). Two canonical signaling pathways are activated through TLR4. The TLR4 adapter protein TIRAP engages the MyD88-dependent signaling pathway that culminates in NF-κB nuclear translocation and pro-inflammatory cytokine production. TRAM, the alternate TLR4 adapter protein, engages TRIF resulting in IRF3 translocation to the nucleus and stimulation of type I interferon response (1).

It is also widely accepted that TLR4 is activated by other agonists including damage-associated molecular patterns, viral proteins and transition metals (4–8). TLR4 was found to mediate immune hypersensitivity reactions to the Group 9/10 transition metals nickel, cobalt and palladium (9–11). Mechanistically, metal binding to TLR4 induces receptor dimerization independent of CD14 and MD-2 (10). Platinum is a Group 10 transition metal that shares chemical properties with nickel and palladium but it is unknown whether it can activate TLR4.

Cisplatin or cis-diamminedichloroplatinum(II), is a platinum-based, highly effective chemotherapeutic frequently used to treat solid tumours in children. In adults it is used to treat ovarian, testicular, cervical, lung, head and neck, and bladder cancers (12). Cisplatin-containing regimens contribute to a 5-year survival rate that approaches 80% in childhood cancer patients and has become an asset to cancer therapy (13). The antitumour activity of cisplatin is based on its formation of intra-strand and inter-strand guanine crosslinks in DNA that prevent the strands from separating, or it alkylates DNA bases causing DNA miscoding (14). This DNA modification activates multiple signal transduction pathways leading to cell-cycle arrest and programmed cell death (15–17).

Despite its effectiveness, cisplatin use is limited by the development of several toxicities that include nephrotoxicity, neurotoxicity and ototoxicity. Although nephrotoxicity can be reversed by saline hydration and mannitol diuresis there is no treatment for cisplatin-induced neurotoxicity or ototoxicity (18). The ototoxic effect of cisplatin leads to permanent bilateral hearing loss and is estimated to affect 26-90% of children treated with cisplatin where age, treatment regimen and concomitant factors also influence susceptibility (19–23). Cisplatin-induced ototoxicity (CIO) can have significant life-long consequences in children by impairing speech and language development, impairing social-emotional development and increasing the risk of learning difficulties (24, 25). Moreover, the likelihood of developing ototoxicity increases in a dose-dependent manner, with nearly 100% of patients receiving high dosages of cisplatin (150-225 mg/m^2^) showing some degree of ototoxicity. This compromises anticancer treatment, potentially impacting overall survival as cisplatin dose reduction or discontinuation is required to mitigate this ototoxicity (26, 27).

CIO is perhaps exacerbated because cisplatin accumulates preferentially in the cochlea of the inner ear (28), and more particularly in the outer hair cells of the Organ of Corti, which are terminally differentiated mechanotransducers and the site of the first steps in sound perception (29). The cochlea is considered a closed system due to its isolated anatomical position and structure and, as such, is not able to rapidly flush out cisplatin and the metabolites generated in response (29). Apoptotic damage in the hair cells of the cochlea is the primary mechanism of cisplatin-induced hearing loss (30).

In the current study, we sought a mechanistic understanding of the signaling pathway activated by cisplatin to enable mitigation of its adverse long-term effects. We found that cisplatin activates TLR4, independently of CD14/MD-2 co-receptors, in a manner reminiscent of nickel. Further, deletion of Tlr4 in a murine inner ear cell line reduced cisplatin-induced ototoxicity. Similarly, knockdown of Tlr4 homologs in zebrafish protected against cisplatin-induced hair cell death. Moreover, we attenuated cisplatin ototoxic responses with the TLR4 chemical inhibitor, TAK-242. These findings provide key insights into the etiology of cisplatin-induced ototoxicity and are crucial to developing protective therapies against CIO, thereby improving the prognosis and long-term health outcomes of cancer patients.

## Results

### Platinum and cisplatin activate TLR4 in vitro

Nickel, palladium and cobalt (Group 9 and 10 transition metals) have been well characterized as TLR4 ligands that induce contact hypersensitivity (9). Given that platinum is a Group 10 transition metal we were interested in determining whether it also could serve as a TLR4 ligand. We investigated this using reporter cell lines that did (HEK-hTLR4) or did not (HEK-null2) stably express human TLR4 and its MD-2/CD14 coreceptors. These isogenic Human Embryonic Kidney (HEK) cell lines also express a reporter of NF-κB activation, where NF-κB induces transcription of secreted alkaline phosphatase (SEAP) reporter; these cells have been used previously to identify TLR4 ligands (11).

We treated HEK-hTLR4 and HEK-null2 cells with platinum (both platinum (II) and (IV) as chloride salts), or LPS or nickel as positive control TLR4 agonists, and monitored NF-κB activation. As expected, we saw a significant > 2-fold activation of NF-κB in HEK-hTLR4 cells treated with LPS or nickel compared to media-only controls (**Fig. 1A**). By contrast, HEK-null2 cells showed no significant change in NF-κB activation, demonstrating the effects were dependent on TLR4. HEK-hTLR4 cells treated with platinum(II) or platinum(IV) also showed significant induction of SEAP activity compared to HEK-null2 cells but the NF-κB activation was less than 2-fold (**Fig. 1A**). Separately, we assessed TLR4 activity by measuring its downstream induction of IL-8 cytokine secretion in the same cells. IL-8 secretion increased by about two orders of magnitude in the HEK-hTLR4 cells for all of the tested agonists (**Fig. 1B**), but not when hTLR4 was absent.

**Figure 1.**
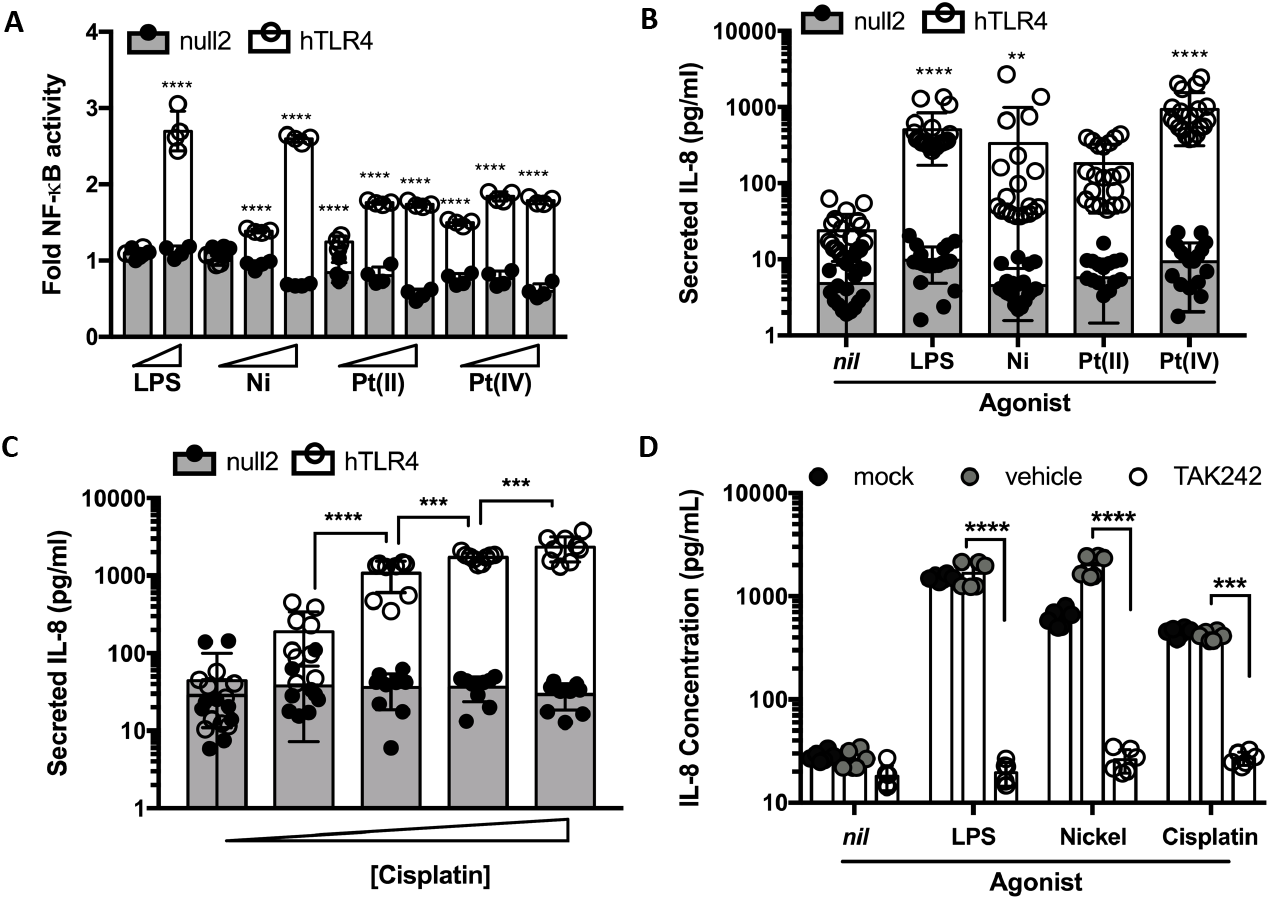
Platinum and cisplatin activate TLR4 *in vitro*. (**A**) Cells expressing TLR4 induced NF-κB activity in response to platinum treatment similarly to known TLR4 agonists. Human embryonic kidney cells that do (HEK-hTLR4), or do not (HEK-null2) stably express *TLR4* were stimulated with LPS (100pg/mL or 1 ng/mL), nickel chloride (100, 200 or 400 μM), platinum (II) chloride and platinum (IV) chloride (25, 50 or 100 μM). NF-κB activity was monitored as a metric of TLR4 activation (normalized to vehicle) (n=4). (**B**) As per panel (A) but secreted IL-8 was monitored as a metric of TLR4 activation (n=20) upon stimulation with LPS (50 pg/mL), nickel (200 μM) and platinum (II and IV) (100 μM). (**C**) TLR4 was sufficient to render cells sensitive to cisplatin in a dose-dependent manner. HEK-hTLR4 and HEK-null2 cells were stimulated with cisplatin (0, 6.25, 12.5, 25 or 50 μM) for 48 hours and IL-8 was quantified in culture supernatants (n=9). (**D**) Cisplatin-induced cytokine secretion, similar to known TLR4 agonists, can be prevented by a TLR4 inhibitor. HEK-hTLR4 cells were pretreated with 4 μM TAK242 (TLR4 inhibitor) or vehicle and then stimulated with cisplatin (25 μM), LPS (50 pg/mL) or nickel chloride (200 μM) (n=4). For all panels data are presented with mean and standard deviation indicated. Data are from 2 (A) or 3 (B and C) independent experiments. Representative data shown in panel D of 2 independent experiments. Statistical analyses was assessed by 2-way ANOVA: A) hTLR4 compared to null2 cells; B) hTLR4 agonist treatment compared to non-treated; C) hTLR4 with comparisons between successive concentrations; and D) comparisons between vehicle and TAK242 treatments. ns, not significant; *, *P*<.05; **, *P*<.01; ***, *P*<.001; ****, *P*<.0001. Multiple comparisons performed using Bonferroni (A, C) or Dunnett’s (B, D) tests.

We next tested whether cisplatin, a platinum-based chemotherapeutic, could also activate TLR4 considering its highly similar composition to platinum chloride. We found that cisplatin also induced IL-8 secretion in HEK-hTLR4 cells more than 100-fold, but not HEK-null2 cells. Furthermore, cisplatin activation of TLR4 was dose-dependent up to 50μM, with pronounced toxicity limiting assessments at higher concentrations (**Fig. 1C**). HEK-null2 cells remained largely unresponsive at higher cisplatin concentrations.

To independently assess the requirement for TLR4 in cisplatin-induced IL-8 secretion in HEK-hTLR4 cells, we repeated our treatments in the presence of a small molecule TLR4 inhibitor (TAK-242) that binds to the intracellular domain of TLR4, disrupting its interactions with cytosolic adaptor proteins (36). Chemical inhibition with TAK-242 mitigated the effect of cisplatin on TLR4 activation similarly to nickel and LPS in HEK-hTLR4 cells (**Fig. 1D**). Taken together, these data demonstrate that platinum and cisplatin behave similarly to nickel and LPS with respect to their ability to activate TLR4.

### Cisplatin activation of TLR4 does not require its co-receptors, MD-2/CD14

We next sought to appreciate the mechanisms of TLR4 activation upon cisplatin activation, as this would influence potential therapeutic design. Canonical TLR4 signaling after binding LPS requires the TLR4 co-receptors MD-2 and CD14, whereas their requirements for metal-based activation of TLR4 are less well-defined (10, 11,37). To examine the role of MD-2/CD14 co-receptors in TLR4 activation we used the HEK-null2 cell line, which lacks TLR4, MD-2 and CD14. HEK-null2 cells were transfected with a human *TLR4* expressing plasmid or empty vector control and assayed for IL-8 secretion upon treatment with TLR4 agonists. As expected, transfection of *TLR4* did not yield a significant increase in secreted IL-8 upon LPS treatment, relative to untreated cells, unless co-transfected with *MD-2* (**Fig. 2A**). By contrast, both cisplatin and nickel significantly enhanced IL-8 secretion in *hTLR4* transfected, but not empty vector transfected HEK-null2 cells. These data suggest that like nickel, cisplatin activation of TLR4 is independent of MD-2/CD14.

**Figure 2.**
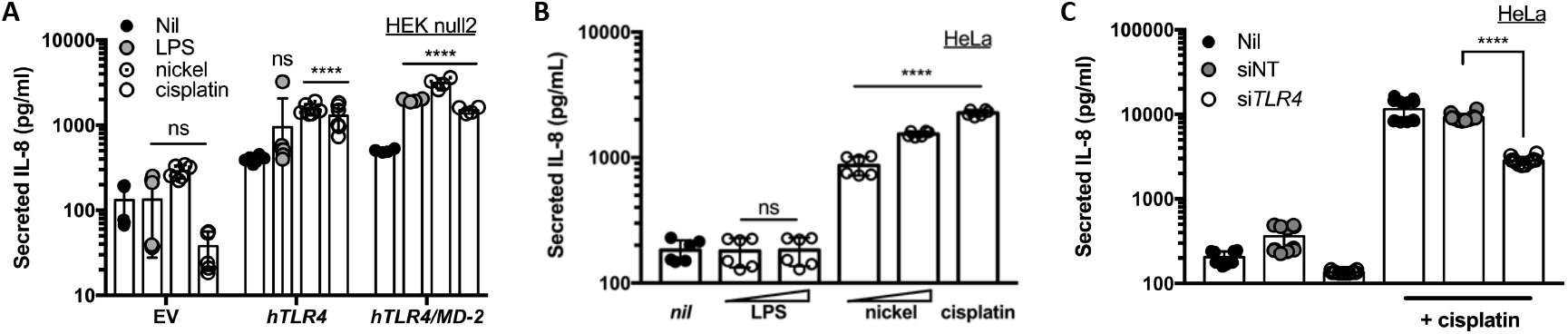
Cisplatin activation of TLR4 is independent of the TLR4 co-receptor, MD-2. (**A**) Transfection of *TLR4* is sufficient for its activation by cisplatin and nickel. HEK-null2 cells were transfected with empty vector (EV), *TLR4* (*hTLR4*), or TLR4 and MD-2 (h*TLR4/MD-2*) and stimulated with LPS (50 pg/mL), nickel chloride (200 μM), or cisplatin (25 μM) (n=6). (**B**) An MD-2-deficient cell line secretes IL-8 cytokine in response to cisplatin and nickel but not LPS. HeLa cells were treated with LPS (10 or 100 ng/mL), nickel chloride (0.4 or 1 mM) or cisplatin (25 μM) for 48 hours. (n=6) (**C**) IL-8 cytokine secretion is dependent on TLR4. HeLa cells were transfected with non-targeting (siNT) or *TLR4*-targeting (si*TLR4*) siRNA constructs and treated with cisplatin (30 μM) for 72 hours. Expression was normalized to untransfected (*nil*) cells (n=12). For all panels secreted IL-8 was quantified as a metric of TLR4 activation and mean and standard deviation are indicated. Data are from two independent experiments (B, C) or representative of 2 independent experiments (A). Statistical analyses were determined in comparison to control treatments: *nil* (A,B) and siNT (C) using 2-way ANOVA (A) or one-way ANOVA (B, C). ns, not significant; *, *P*<.05; **, *P*<.01; ***, *P*<.001; ****, *P*<.0001. Multiple comparisons performed using Dunnett’s tests.

To further investigate the requirement of MD-2/CD14 co-receptors for cisplatin activation of TLR4 we used HeLa cells that have been reported to lack MD-2 expression (38). We treated HeLa cells with cisplatin, nickel and LPS and observed that cisplatin and nickel, but not LPS, induced significant IL-8 secretion (**Fig. 2B**). To confirm that IL-8 secretion elicited by cisplatin in HeLa cells was dependent on TLR4, we repeated cisplatin treatments in HeLa cells transfected with non-targeting or *TLR4*-targeting siRNA. We determined that siRNA treatment reduced TLR4 expression by >75% (**Fig. S1**). Following *TLR4* knockdown we observed 70% lower cisplatin-induced IL-8 secretion indicating that secretion of this cytokine is mediated by TLR4 (**Fig. 2C**). Taken together these data indicate that TLR4 co-receptors are dispensable for cisplatin activation of TLR4.

### Tlr4 deletion mitigates cisplatin ototoxic responses in a murine inner ear cell line

Having shown that cisplatin can act as an agonist of TLR4 to induce a pro-inflammatory response *in vitro*, we next asked whether TLR4 plays a role in mediating the molecular events that contribute to cisplatin-induced ototoxicity. Cisplatin treatment induces the generation of reactive oxygen species (ROS) in the cochlea, which appear to be critical mediators of CIO (39, 40). Hallmarks of *in vitro* cisplatin ototoxic responses, as modelled by Organ of Corti cell lines, include increased pro-inflammatory IL-6 signaling, which can upregulate ROS generation that in turn influence morphological and functional alterations leading to apoptotic cell death (41–43).

We used the mouse inner ear Organ of Corti cell line HEI-OC1, which provides a popular *in vitro* model of drug-induced hearing loss (44), and mutated *Tlr4* by CRISPR/Cas9. We established single-cell clones of *Tlr4*-edited cells, along with nontargeting guide RNA-edited control cells and conducted a primary screen to identify clones with diminished LPS responses. Sanger sequencing at the *Tlr4* locus identified a clone with frame-shift mutations in exon 1 (one adenine insertion or a four nucleotide deletion; **Fig. S2A**). Compared to control cells, the deletion clone exhibited decreased Tlr4 protein abundance (**Fig. S2B**), significantly reduced binding/internalization of a fluorescent LPS analog (**Fig. S2C**) and significantly reduced LPS-induced cytokine secretion (**Fig. S2D**). Importantly, LPS-induced IL-6 secretion was enhanced 4-fold upon complementation with ectopically expressed *Tlr4* in the deletion cells, compared to less than 2-fold in control cells (**Fig. S2D**, ***inset***). Taken together with the genetic data, these results confirm a *Tlr4* deletion in CRISPR targeted HEI-OC1 cells.

To examine the impact of the *Tlr4* deletion on cisplatin ototoxic responses, we treated *Tlr4* deletion and control HEI-OC1 cells with cisplatin for 24 hours to measure apoptosis, proinflammatory cytokine secretion and intracellular ROS generation. With increasing cisplatin concentrations, *Tlr4* deletion cells showed less apoptotic and concomitantly more live cells, compared to control cells (**Fig. 3A**). Similarly, we observed a significant decrease in cisplatin-induced ROS formation in *Tlr4* deletion cells at higher cisplatin concentrations (**Fig. 3B**). Moreover, *Tlr4* deleted cells had reduced IL-6 secretion in response to cisplatin treatment compared to control cells (**Fig. 3C**). Taken together, these data indicate that TLR4 is an important mediator of cisplatin ototoxic responses in inner ear hair cells.

**Figure 3.**
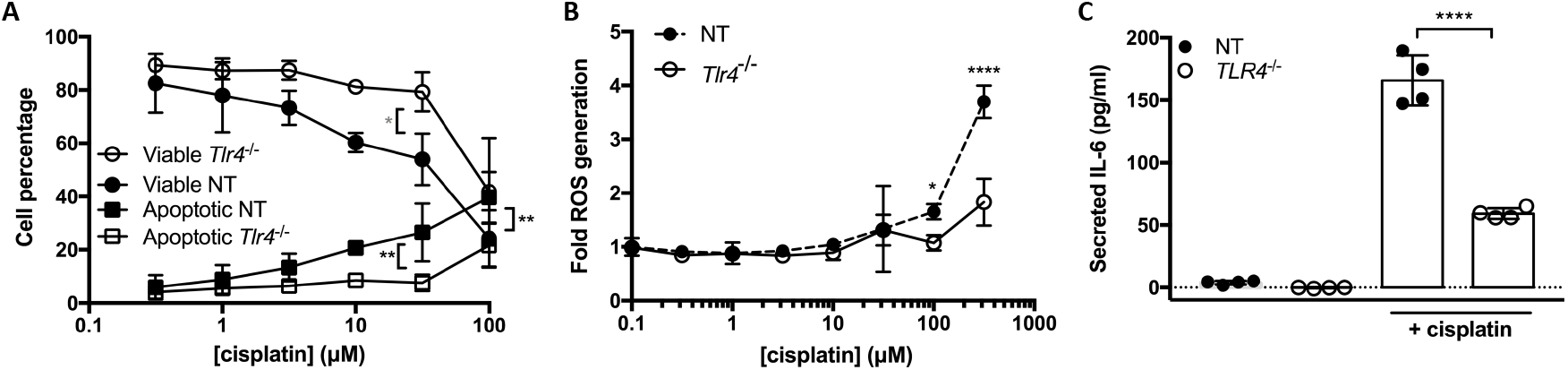
Deletion of *Tlr4* in a murine ear outer hair cell line (HEI-OC1) reduces cisplatin-induced ototoxic responses. *Tlr4* deletion (*Tlr4*^-/-^) increased cell viability and inhibited apoptosis induction (**A**, n=3), diminished ROS generation (**B**, n=4), and reduced IL-6 secretion (**C**, n=4), compared to non-targeting (NT) control cells. Cells were analyzed by flow cytometry (AnnexinV/PI; **A** and DCFH-DA; **B**) or ELISA (**C**). For all panels data are presented with mean and standard deviation indicated. Data are from 3 (A) or 2 (B and C) independent experiments. Statistical comparisons to NT at the same cisplatin concentration were assessed by 2-way ANOVA (A, B) or one-way ANOVA (C). *, *P*<.05; **, *P*<.01; ***, *P*<.001; ****, *P*<.0001. Multiple comparisons performed using Bonferroni tests. Note for (A) grey asterisk refers to viability comparisons.

### Cisplatin-induced toxicity is consistent with its primary activation of TLR4

It has been previously reported that cisplatin induces the expression of *Tlr4* leading to subsequent activation by LPS to potentiate cisplatin ototoxicity (45). This model describes a secondary effect of cisplatin on TLR4 activation. While our data demonstrating that cisplatin toxicity responses depend, at least in part, on Tlr4 could be consistent with this model, our observations in the HEK-hTLR4 system suggest that cisplatin has a primary effect on TLR4 activation (e.g. co-receptor-independent TLR4 activation). To further characterize the effect of cisplatin in an ear outer hair cell line we conducted kinetic analyses of Tlr4 activation to distinguish between primary (early) and secondary (later) effects. We examined IL-6 secretion over time in HEI-OC1 cells stimulated by the TLR4 agonists, cisplatin and LPS. We observed that cisplatin- and LPS-induced IL-6 secretion followed similar kinetics for 4 hours with cisplatin continuing to induce secretion after 24 hours, unlike LPS (**Fig. 4A**). Cisplatin and LPS co-treatment had an additive effect on IL-6 secretion starting at 2 hours post-treatment (**Fig. 4A**). These data suggest that cisplatin is activating TLR4 in a primary manner.

**Figure 4.**
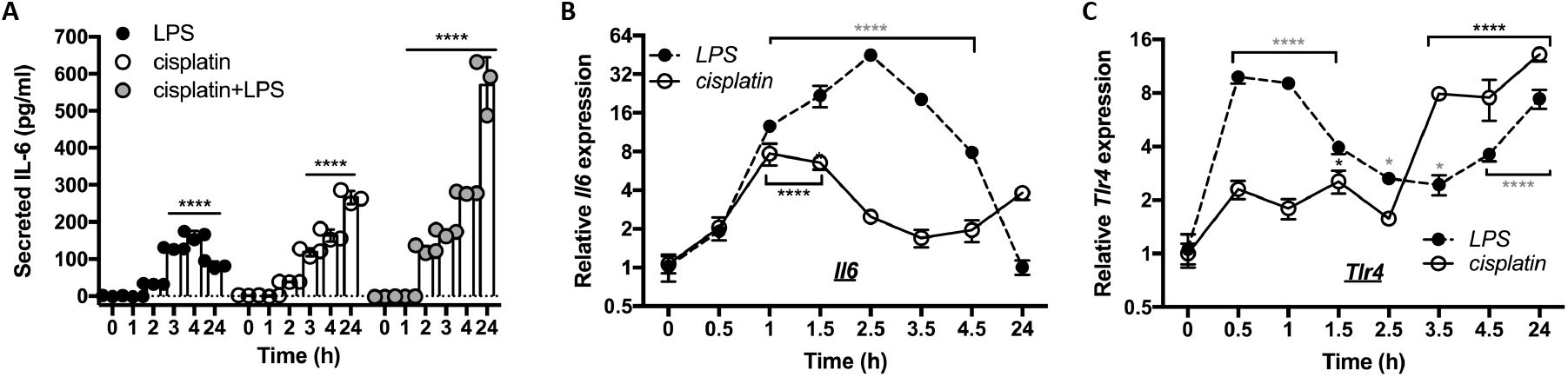
Cisplatin has a primary role in Tlr4 activation in HEI-OC1 cells. (**A**) Cisplatin and LPS elicit IL-6 secretion with similar kinetics at early time points. HEI-OC1 cells were treated with LPS (100 pg/mL), cisplatin (20μM) or both and secreted IL-6 was quantified as a metric for TLR4 activation (n=3). Data (mean ± SD) are representative of 3 independent experiments. (**B**, **C**) Cisplatin differentially induces *Il6* and *Tlr4* expression in comparison to LPS. HEI-OC1 cells were treated with cisplatin (20μM) or LPS (1 ng/mL) and *Il6* (B) or *Tlr4* (C) transcript levels were quantified at the indicated time points (n=4). Mean and standard deviation are indicated. Data are from 2 independent experiments (B,C) or representative of 3 independent experiments (A). *, *P*<.05; ****, *P*<.0001 as determined by 2-way ANOVA with Dunnett’s multiple comparison test to 0 hr time point. Note for (B and C) grey asterisks refer to LPS comparisons.

To further investigate TLR4 activation by cisplatin we treated HEI-OC1 cells with cisplatin or LPS and total RNA was extracted. We quantified the relative expression of both *Il6* and *Tlr4* over time in response to cisplatin and LPS treatment. LPS treatment caused *Il6* expression to rise sharply after 1 hour, peaking at 2.5 hours and returning to basal levels after 24 hours (**Fig. 4B**). Cisplatin treatment also showed *Il6* expression peaking after 1 hour, albeit at half the expression level as LPS treatment. Notably, *Il6* expression was elevated after 24 hours in response to cisplatin treatment (**Fig. 4B**), which correlates with the IL-6 secretion kinetics (**Fig. 4A**). Interestingly, there was a notable disparity in *Tlr4* expression patterns following cisplatin and LPS treatments. LPS treatment induced *Tlr4* expression after only 30 minutes, followed by a gradual reduction until peaking again after 24 hours (**Fig. 4C**). By contrast, cisplatin treatment caused *Tlr4* expression to remain relatively stable until sharply rising after 3.5 hours (**Fig. 4C**). The kinetics of cisplatin-induced cytokine secretion in our experiments, coupled with our observations that cytokine gene expression preceded *Tlr4* expression in response to cisplatin treatment, supports a model where cisplatin has a primary effect on Tlr4 activation.

### Zebrafish homologs of TLR4 are required for cisplatin-induced ototoxicity

Having shown that TLR4 played a critical role mediating cisplatin ototoxicity responses *in vitro* we sought to examine the role of TLR4 in an *in vivo* CIO model. We chose to use zebrafish because it is a robust and widely-accepted model of ototoxicity (46–52). Using established assays, we scored the health of neuromasts, which are mechanotransducing hair cells that bear structural, cellular, and physiological similarities to Organ of Corti outer hair cells (53). Neuromast health can be visualized using the fluorescent dye DASPEI, which accumulates and stains viable hair cells. We used a dose-response format to establish a dose of cisplatin that robustly reduced hair cell viability in our hands (**Fig. 5A**). A concentration of 15μM was chosen for subsequent experiments.

**Figure 5.**
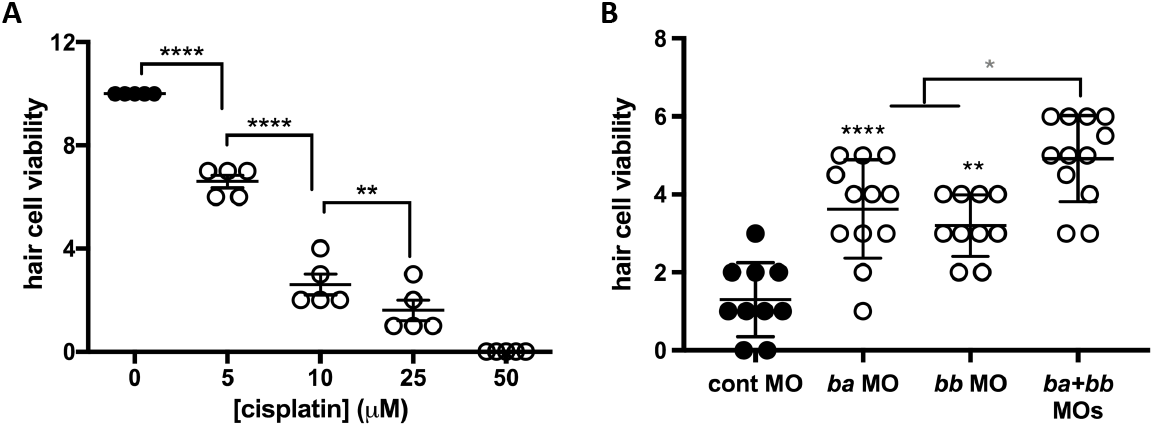
Zebrafish Tlr4 mediates cisplatin-induced otoxicity *in vivo*. (**A**) Larval fish were treated with increasing concentration of cisplatin. Hair cell viability was assessed by DASPEI staining and scored for viability using fluorescence microscopy. Each data point (circles) represents a score of hair cell integrity in an individual animal (taken from multiple samples per animal), whereas lines represent mean ± SD. (**B**) Tlr4 knockdown in zebrafish ameliorates cisplatin ototoxicity. Gene knockdown in larval fish was accomplished by pre-treatment with control-, *tlr4ba*- and/or *tlr4bb*-targeting morpholino oligonucleotides prior to treatment with 15 μM cisplatin. Data are presented as in (A). *, *P*<.05; **, *P*<.01; ****, *P*<.0001 as determined by oneway ANOVA with Tukey multiple comparison testing. Comparisons are to control morpholino in (B) except as indicated (grey asterisk).

Zebrafish have two *tlr4* genes, designated *tlr4ba* and *tlr4bb* that are orphan receptors. They are not activated by LPS but chimeric experiments show that they are linked to the NF-κB signaling pathway (35). Prior to bath application of cisplatin, we knocked down the *tlr4ba* homolog, the *tlr4bb* homolog, or both *tlr4ba* and *tlr4bb* homologs, using morpholinos that were previously validated thoroughly for specificity and efficacy (33–35). Knockdown of either *tlr4ba* or *tlr4bb* was significantly protective against CIO (**Fig. 5B**). Moreover, a protective effect against cisplatin-induced neuromast toxicity was observed with two independent *tlr4bb*-targeting morpholinos that disrupt gene splicing or translation (**Fig. S3**). Notably, we observed that protection from CIO could be further enhanced by combinatorial knockdown of both *tlr4ba* and *tlr4bb*, further supporting the role of zebrafish *tlr4* in CIO (**Fig. 5B**).

### Chemical inhibition of TLR4 decreases cisplatin ototoxicity responses in vitro

Our data suggest TLR4 may be a druggable therapeutic target to mitigate CIO. We next sought to examine the effect of a TLR4 chemical inhibitor on cisplatin toxicity in HEI-OC1 cells. We blocked the TLR4 signaling pathway in these cells by pre-treating them with TAK-242 or vehicle control prior to treatment with LPS or cisplatin and concurrent TAK-242/vehicle treatment. We chose this inhibitor, rather than one that targets TLR4/MD-2 interactions, because our findings indicated that TLR4 activation by cisplatin is independent of MD-2. Analysis of secreted IL-6 levels indicated that cells treated with TAK-242 released significantly less IL-6 protein in comparison to the vehicle control in response to both cisplatin and LPS agonists (**Fig. 6A**). Similarly, ROS generation was significantly reduced in cells treated with cisplatin and TAK-242 compared to cisplatin and vehicle treatments (**Fig. 6B**). Overall, these results indicate that cisplatin ototoxic responses that underlie hearing loss can be mitigated using a chemical inhibitor of TLR4 in an ear outer hair cell line.

**Figure 6.**
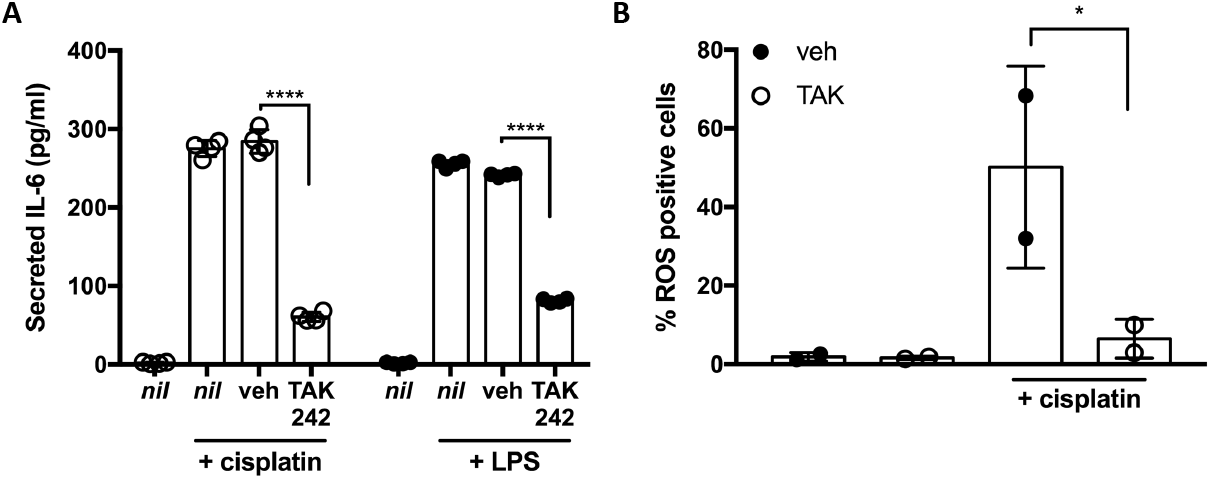
Small molecule inhibition of Tlr4 mitigates cisplatin-induced ototoxic responses in HEI-OC1 cells. Treatment of HEI-OC1 cells with the TLR4 inhibitor, TAK242 significantly reduces cisplatin-induced IL-6 secretion (**A**, n=4) and ROS generation (**B**, n=2). HEI-OC1 cells were pretreated with DMF vehicle (veh), TAK242 (4μM) or left untreated (*nil*) and subsequently treated with LPS (10 ng/mL) or cisplatin (20μM) as indicated. Data, presented with mean ± SD, are from 2 independent experiments. The percent ROS positive cells was determined by flow cytometry using Total ROS-ID reagent. Statistical analyses were compared to vehicle by 2-way ANOVA with Dunnet’s multiple comparison testing (A) or one-way ANOVA with Tukey multiple comparison testing (B). *, *P*<.05; **, *P*<.01; ****, *P*<.0001.

## Discussion

In this work, we have shown that TLR4 is a critical mediator of cisplatin-induced ototoxic responses in an ear outer hair cell line and in zebrafish. This is the result of cisplatin activating TLR4 based on its structural similarity to platinum chloride. Moreover, we show for the first time that a small molecule inhibitor of TLR4 can mitigate cisplatin ototoxic responses in ear outer hair cells. Taken together, this work sets the stage to focus on TLR4 as a therapeutic target for the mitigation of CIO and establishes appropriate model systems for these preclinical efforts.

ROS generation and apoptosis induction in outer hair cells are considered the basis of how cisplatin kills these critical mechanotransducing cells; however, it is poorly understood how these responses are elicited. Previous studies have identified key roles for the TNFa and NF-κB inflammatory signaling pathways in transducing cisplatin ototoxic responses but the upstream processes were less defined (43, 54). Our work now identifies TLR4 as a bridge between cisplatin and these signaling pathways since TLR4 activation can induce NF-κB signaling and TNFa secretion (1,55). Interestingly, some reports have demonstrated the activation of TLR4 as one of the main pathways causing ototoxicity by aminoglycoside treatment and cochlear inflammation after acoustic injury (56–58). The specific ligands that activate TLR4 in these conditions are not defined; however, in general, each of these types of damage increases *Tlr4* expression levels in the cochlea within hours (56, 58).

It has been reported that a *Tlr4* deletion in C3H/HeJ mice partially mitigated cisplatin-induced nephrotoxicity (59). This work was interpreted to suggest, but did not confirm, that damage-associated molecular patterns activate TLR4 in C3H/HeJ mice and cause cisplatin-induced renal toxicity (59). Others have reported that LPS acts coordinately with cisplatin to increase inflammatory responses and cellular damage in renal and cochlear cells (45, 60). Specifically, Oh et al. suggested that cisplatin plays a secondary role in TLR4 activation by upregulating the expression of *Tlr4* for subsequent activation by LPS (45). Our study contributes to a new understanding of the association of cisplatin and TLR4 in the induction of CIO. Our data clearly indicate that cisplatin can activate TLR4 *in vitro* and does so in a manner that is mechanistically disparate from LPS. We observed that TLR4 activation by cisplatin in an ear outer hair cell line (assessed by cytokine secretion) occurred with similar kinetics to TLR4 activation by LPS. Furthermore, we noted that cisplatin treatment induced *Tlr4* expression at later time points than LPS. Taken together, our data argue that cisplatin has a primary effect on TLR4 activation, similar to the TLR4 agonists LPS and nickel.

While highly structurally distinct from LPS, metal contact allergens have been shown to signal through direct TLR4 interactions (9, 11). Our results suggest that platinum chloride and cisplatin are also capable of activating TLR4. Given that platinum is also a Group 10 transition metal, we speculated that platinum and cisplatin may activate TLR4 in a manner analogous to nickel, rather than LPS. In line with this, our *in vitro* analyses showed that unlike LPS, cisplatin was able to activate TLR4 signaling in the absence of the TLR4 co-receptors MD-2 and CD14, as we observed with nickel. Interestingly, although we observed that nickel was able to signal through TLR4 without MD-2, other studies have found that this co-receptor is required for effective signaling through human TLR4 (61).

Nickel is proposed to form critical interactions with TLR4 histidine residues (456 and 458) on the ectodomain of human TLR4 that facilitate the dimerization of TLR4 complexes and subsequent signaling(10, 11). The role of these residues in cisplatin activation of TLR4 remains to be studied and could help glean information on whether cisplatin behaves as a direct ligand of TLR4 by analogy to nickel. While our kinetic analyses of cisplatin responses in HEI-OC1 cells strongly suggest that cisplatin has a primary effect on TLR4 activation, it does not confirm a direct interaction between cisplatin and TLR4 and further study is required to establish this.

Zebrafish have proved to be a robust model system for studying cisplatin-mediated hair cell death by monitoring neuromast viability (32, 48, 62). Moreover, this model has been used to investigate potential otoprotective therapies (46, 51, 63–69). It is however, well recognized that zebrafish tlr4ba/bb are distinct from mammalian TLR4. *TLR4* homologs appear to have been lost from the genomes of many fish species, suggesting a very disparate role for TLR4 compared to its centrality in mammalian responses to LPS endotoxin. Indeed, zebrafish *TLR4* homologs appear to be unresponsive to LPS, which has been attributed to a lack of an MD-2 ortholog in zebrafish (35). Nevertheless, chimeric mammalian TLR4 and zebrafish tlr4ba/bb constructs studied *in vitro* showed that the intracellular domains of the zebrafish proteins could interact with downstream signaling components in the TLR4 pathway (35). Our finding that zebrafish TLR4 homologs are required for CIO is consistent with their expression in zebrafish hair cells (70, 71) and is in line with the *in vitro* and murine inner ear cell studies presented in this work. This further supports the identification of TLR4 as a key mediator of cisplatin-induced ototoxicity. These results suggest that zebrafish can be an important model for screening TLR4 antagonists as putative otoprotectants against CIO. They also raise the intriguing possibility that zebrafish tlr4ba/bb, heretofore orphan receptors with no known agonist, could be sensors of Group 10 transition metals.

In aggregate, our data argue that cisplatin plays a primary role in activating TLR4, independently of LPS, suggesting that TLR4 is a critical mediator of CIO. This is reinforced by our observation that genetic or chemical inhibition of Tlr4 in an ear outer hair cell line or in zebrafish, significantly reduced cisplatin toxicity. To our knowledge, our TAK242 data represents the first demonstration that a small molecule inhibitor of TLR4 can mitigate cisplatin toxicity. Given that cisplatin activation of TLR4 is distinct from LPS, this affords an opportunity to develop tailored therapies that specifically target cisplatin-activation of TLR4. The findings in this study bring us closer to understanding the mechanisms involved in CIO, with TLR4 as a primary target for the rational design of otoprotectants, and bettering health outcomes for cancer patients while conserving the success of cisplatin chemotherapy.

## Materials and Methods

### Cell culture and treatments

The murine inner ear cell line HEI-OC1 (a kind gift from Dr. Federico Kalinec, UCLA) were grown in DMEM supplemented with 10% FBS (Gibco, 123483-020), 5% penicillinstreptomycin (1 unit penicillin/mL and 0.1 mg streptomycin/mL, Sigma, P4333). HEI-OC1 cells were grown at 33°C in the presence of 10% CO_2_. HEK (Human Embryonic Kidney) -hTLR4, and -null2 cell lines (cat# hkb-htlr4 and hkb-null2, Invivogen) are isogenic reporter cell lines stably transfected with a secreted alkaline phosphatase reporter under the control of five tandem NF-κB response elements. HEK-hTLR4 cells also stably express human *TLR4, CD14* and *MD-2*. These cells were grown in DMEM supplemented with 10% FBS, 5% penicillin-streptomycin, and 100μg/mL Normocin (Invivogen) at 37°C and 5% CO_2_. Cells were routinely seeded in 96-well plates (5 x 10^3^ cells/well), 24-well plates (7.0 x 10^4^ cells/well), 12-well plates (1.1 x 10^5^ cells/well) or 6-well plates (1.5-2.5 x 10^5^ cells/well), Cisplatin (Teva, 02402188), LPS (Invitrogen, L23351), nickel chloride hexahydrate (Sigma, 654507), platinum (II) chloride (Sigma, 520632) or platinum (IV) chloride (Sigma, 379840) was added to cells 48 hours after seeding in fresh media. Vehicle (DMF; Fisher Scientific, D1331) or TAK242 (Cayman, 243984-11-4) was added to cell culture in fresh media 1 hour prior to treatments. Following 1 hour pre-treatment, media was aspirated and vehicle or TAK242 were added to cells in combination with cisplatin, LPS, or nickel treatments for 24 or 48 hours. All reagents were assessed for endotoxin contamination ≥0.125 εU using Pyrotell Gel Clot Formulation kit for bacterial endotoxin testing (Pyrotell, GS125-5). LPS and low endotoxin water (<0.005 εU; HyClone, SH30529.02) were used as positive and negative controls, respectively. All TLR4 agonists (except LPS) tested negative for endotoxin.

### CRISPR-Cas9 mediated Tlr4 knock out

Mouse *Tlr4* was targeted for mutation in HEI-OC1 cells using TrueCut™ Cas9 Protein V2 (Invitrogen, A36498), TrueGuide™ tracrRNA (Invitrogen, A35507) and TrueGuide™ Syn crRNA (Invitrogen, A35509-CRISPR511653) and gRNA (Target: GATTCAAGCTTCCTGGTGTC). TrueGuide™ Syn crRNA, Negative Control, (Invitrogen, A35519) was used as a non-targeting crRNA in this assay. Gene editing efficiency was verified using the GeneArt™ Genomic Cleavage Detection kit (Invitrogen, A24372) in a pooled cell population. Procedures were carried out based on the manufacturer’s protocols. Single-cell clones were then isolated for further validation using limited dilution in 96-well plates. *Tlr4* deletion clones were then screened for loss of LPS-induced IL-6 cytokine secretion. Genomic DNA from selected clones was amplified at the *Tlr4* locus and analyzed by Sanger Sequencing using primer pair: 5’-CCTCCAGTCGGTCAGCAAAC-3’ and 5-’CTAAGCAGAGCACACACAGGG-3’.

### Immunocytochemistry

Immunofluorescence staining was used to examine levels of TLR4 protein from control and *Tlr4*-deleted HEI-OC1 cells. Cells were seeded in a 96-well plate and fixed after 24 hours in 4% paraformaldehyde for 20 minutes at room temperature. After washing 3X with PBS, blocking was performed using 3% BSA for 1 hour. Cells were then stained using anti-mouse TLR4 primary antibody (Invitrogen, 13-9041-80) and Alexa-488 Fluor Goat anti-mouse IgG secondary antibody (Jackson Immunoresearch; 115-545-146). Antibodies were used at 1:200 dilution. Finally, secondary antibody was removed by washing with PBS 3X and PBS was added to the wells. Images were acquired using an Evos FL Auto microscope and manufacturer’s software.

### Cell viability assays

MTT reagent (ACROS, 158990010) was added to 1 mg/mL to seeded cells, 24 or 48 hours post-treatment. When required, aliquots of the supernatant were collected for ELISAs before the addition of MTT. Plates were incubated at 33°C at 10% CO_2_ (HEI-OC1) or 37°C at 5% CO_2_ (HEK) for 4 hours in the dark. Next, supernatants were replaced with DMSO (Sigma, D109), and incubated with shaking at room temperature for 20 minutes. Absorbance at 590nm was collected in a plate reader (SpectraMax i3x, Molecular Devices). For the purposes of cell viability dose-response curves, the mean absorbance for a no-treatment control was considered 100% cell viability so that cell viability (%) of treatment = (absorbance_treatment_/absorbance_control_) x 100.

### NF-κB activation assays

To measure TLR4 activation in the HEK-null2 and HEK-hTLR4 cells, an integrated NF-κB reporter system was used. The reporter is a secreted alkaline phosphatase enzyme that is transcriptionally regulated by NF-κB. The secreted alkaline phosphatase assay was performed according to the manufacturer’s instructions with slight modifications. Cells were cultured in DMEM containing 10% FBS, 1% Penicillin/Streptomycin and 0.1mg/mL Normocin (ant-nr-2, Invivogen). HEK-Blue selection (hb-sel Invivogen) and zeocin (ant-zn-05 Invivogen) were applied to HEK-hTLR4 and HEK-null2 cells, respectively, every 5 passages to maintain stable cell transfection. Cells were seeded in a 96 well dish at 1.4×10^5^ cells/mL (HEK-hTLR4) and 2.8×10^5^ cells/mL (HEK-null2) in HEK detection media (Cat# hb-det3 Invivogen) as per manufacturer’s protocol. Cells were treated upon seeding with media, DMF (vehicle for platinum (II) chloride) or TLR4 agonists. Alkaline phosphatase activity was measured by reading absorbance at 620 nm after 36 hours of stimulation (SpectraMax i3x reader, Molecular Devices).

### ELISA assays

As an alternate method of assessing TLR4 activation, IL-6 and IL-8 secretion was quantified in HEI-OC1 and HEK cells, respectively. Colorimetric protein assays were conducted using commercial human IL-8 ELISA and mouse IL-6 ELISA kits (Invitrogen; 88-8086, 88-7064) according to the manufacturer’s protocol. Supernatants were collected from 12 well plates or 24 well plates 0, 0.25, 0.5, 1, 2, 3, 24 or 48 hours posttreatment (leaving half the volume in the well for subsequent MTT assays). Protein secretion was normalized to the number of viable cells to account for agonist toxicity.

### LPS internalization assay

We assessed the extent of LPS internalization to characterize our HEI-OC1 *Tlr4* deletion cell line compared to the non-targeting control cells. *Tlr4*-deletion and control HEI-OC1 cells were grown up to 90% confluence in 6-well plates in complete DMEM medium. Cells were then treated with 5 μg/mL ultrapure Alexa-488 Fluor™ LPS (Life Technologies, L23351) or with non-fluorescent LPS (eBioscience™, 00-4976-93) as a control for 4 hours. Cellular LPS internalization was assayed via flow cytometry after quenching with 1mg/mL Trypan blue. 50,000 cells were read and gated by forward and side scatter for selecting live cells and then single cells. Cellular uptake of Alexa-488 Fluor™ LPS was shown based on the median fluorescence intensity (MFI) at 488 nm excitation wavelength and 525 nm emission wavelength. Samples treated with non-fluorescent LPS were used to subtract background auto-fluorescence. Data analysis was performed using FlowJo_V10 software.

### Apoptosis assay

We monitored apoptosis induction as a hallmark response of cisplatin treatment *in vitro*. HEI-OC1 cells were treated with increasing concentrations of cisplatin for 24 hours. After 24 hours, cells were harvested and washed once with PBS and once with 1x Annexin V binding buffer (ThermoFisher Scientific, V13246). Following washes, cells were re-suspended in 100 μL Annexin V binding buffer. Cells were then stained following the manufacturers’ protocol. In brief, 5 μL of FITC-Annexin V and 1 μL of diluted propidium iodide solution (100 μg/mL working solution) was added to each sample. Samples were then incubated at room temperature in the dark for 15 minutes. Samples were diluted with 400 μL Annexin V binding buffer and acquired on an Attune NxT flow cytometer (ThermoFisher Scientific). A minimum of 10,000 events were acquired in each sample. Following the acquisition, samples were analyzed using FlowJo (BD Biosciences).

### ROS detection assay

We monitored ROS generation as a hallmark response of cisplatin treatment *in vitro* using two different ROS indicators. 2’,7’-dichlorofluorescein diacetate (DCFH-DA) (Sigma, D6883) was used to measure intracellular ROS levels. Serum-free DMEM containing 4μM DCFH was added to the cells post-treatment in a 96-well plate and incubated at 33°C at 10% CO_2_ for 1 hour. Cells were washed with PBS two times and resuspended in 100 μL of PBS. ROS generation was monitored using a plate reader (SpectraMax i3x) at emission and excitation wavelengths of 538 nm and 485 nm, respectively. Complete media was added to the cells for subsequent MTT analysis for normalization as outlined above. Normalized values were then used to calculate ROS fold induction for each sample. Normalized values of no-treatment cells were used as a baseline value to calculate fold induction [fold of induction of ROS= (fluorescence_sample_/absorbance_sample_)/(fluorescence_no treatment control_/absorbance_no treatment control_].

For total ROS measurements in response to TAK242 treatment, we used the Total ROS-ID detection kit (Enzo Life Sciences, ENZ-51011). Cells were trypsinized and washed with 1X ROS Wash Buffer. Cells were then stained following the manufacturers’ protocol. In brief, cells were re-suspended in 500μl of the ROS Detection Solution for 30 minutes. No washing was required prior to sample analysis using Attune NxT flow cytometer (ThermoFisher Scientific). A minimum of 10,000 events were acquired in each sample. Following acquisition, samples were analyzed using FlowJo (BD Biosciences).

### Cell transfections

HEK-null2 cells were transfected with a human *TLR4* expression clone (pcDNA3-TLR4-YFP was a gift from Doug Golenbock – http://n2t.net/addgene:13018) and human MD-2 expression clone (Origene; RC204686) to assess cytokine secretion in response to TLR4 agonist treatments. HEI-OC1 cells were transfected with a mouse *Tlr4* expression clone (Origene; MR210887) to test for complementation of the *Tlr4* deletion strain. Transfections of HEK cells were carried out with Lipofectamine 3000 (Invitrogen, L30000 IS) and of HEI-OC1 cells with jetPRIME (Polyplus, CA89129-924) reagents in 24 well plates, with 0.5μg of DNA according to the manufacturer’s specification. Cells were treated 24 hours post-transfection with cisplatin, LPS, or nickel.

### qPCR assay

HEI-OC1 cells were seeded in 6-well plates with 250,000 cells/well. Cells were allowed to grow for 48 hours to achieve a 70% confluency prior to treatment with cisplatin. Cells were all treated simultaneously. RNA extraction (BioRad Aurum Total RNA Mini Kit) was performed at the appropriate time points after cisplatin addition (0, 1, 2, 3, 4, and 24 hours post-treatment). cDNA synthesis was performed using the iScript gDNA-Clear cDNA Synthesis kit (BioRad, 1725034). Final qPCR was performed using TLR4 (qMmuCIP0035732) and HPRT (qMmuCEP0054164) primer-probe assays obtained from BioRad and the associated SSO Advanced Universal Probe Super Mix kit (BioRad 1725281).

### siRNA gene knockdown

Cells were seeded in 6-well plates with 150,000 cells/well. Cells were allowed to grow for 24 hours to achieve 50-60% confluency and were transfected with 5nM of Non-Targeting/Negative Control siRNA or the appropriate *TLR4* siRNA (hs.Ri.TLR4.13) using a dsiRNA TriFECTa Kit from Integrated DNA Technologies in conjunction with RNAiMAX transfection reagent (Thermofisher). Cells were allowed to grow for another 24 hours following transfection prior to treatment with cisplatin. Exposure to cisplatin continued for 72 hours prior to the collection of supernatants detection of secreted IL-8 by ELISA and the completion of MTT cell viability assays for normalization as previously indicated.

### Animal ethics and zebrafish husbandry

Zebrafish were kept at the University of Alberta following a 14:10 light/dark cycle at 28°C cycle as previously described(31). They were raised, bred and maintained following an institutional Animal Care and Use Committee approved protocol AUP00000077, operating under guidelines set by the Canadian Council of Animal Care.

### Assessing CIO in larval zebrafish

Wildtype (AB strain) zebrafish were grown to 5 days post fertilization (dpf) in standard E3 embryo media (31) and were bath treated with either 0, 5, 10, 25 or 50μM of cisplatin in 6-well plates, with 10-15 zebrafish larvae per well. After a 20 hour incubation with cisplatin at 28°C, wells were washed with embryo media before the fish were incubated in media containing 0.01% 2-[4-(dimethylamino) styryl]-1-ethylpyridinium iodide (DASPEI, Sigma-Aldrich) to stain for neuromast mitochondrial activity for 20 minutes. Wells were washed again in embryo media and zebrafish larvae anaesthetized with 4% tricaine. Neuromasts were imaged under a Leica M165 FC dissecting microscope equipped with a fluorescent filter. A standard scoring method for zebrafish hair cell viability was used (32): five posterior lateral line (PLL) neuromasts for each fish were assigned a score representing cell viability based on DASPEI fluorescent intensity (2 for no noticeable decline, 1.5 for minor decline, 1 for moderate decline, 0.5 for severe decline and 0 for complete loss of fluorescent intensity). These five scores were summed for each individual (10= all hair cells appear normal and viable; 0=intense ototoxicity).

### Morpholino knockdown of TLR4 homologs

Previously validated anti-sense knockdown reagents (Morpholinos (33–35)) against *tlr4ba* and *tlr4bb* (Gene Tools, LLC; Philomath, OR) were delivered to developing zebrafish. Two *tlr4bb* morpholinos were used, the first translation blocking: *trl4bb*-MO1 (5’-AATCATCCGTTCCCCATTTGACATG-3’) the second splice blocking: *tlr4bb*-MO2 (5’-CTATGTAATGTTCTTACCTCGGTAC-3’). A splice blocking *tlr4ba*-MO2 (5’-GTAATGGCATTACTTACCTTGACAG-3’) was also used. All gene-specific morpholinos have been previously described and thoroughly vetted for efficacy and specificity to the gene target (33–35). A standard control morpholino (5’-CCTCTTACCTCAGTTACAATTTATA-3’) was used as a negative control. Injection solution for morpholinos consisted of 0.1M KCl, 0.25% dextran red, either the standard control or gene-specific morpholinos to effective dose and nuclease-free water. One-cell stage newly fertilized embryos were positioned on an agarose plate and injected with 5ng of morpholino. At 2dpf gene-specific morpholino injected fish, control morpholino injected fish and un-injected fish were added to separate wells of a 6-well plate with 10-15 fish per well. Fish were incubated with 15μM cisplatin for 20-hours before being washed, DASPEI stained, imaged and analyzed as described above.

### Statistical analyses

TLR4 activation across multiple cell lines was analyzed by 2-way ANOVA with Bonferroni multiple comparisons test between samples and Dunnett’s multiple comparisons test against a control sample (*nil* or vehicle). TLR4 activation in a single cell line was analyzed by one-way ANOVA using Dunnett’s multiple comparisons test to a control sample (*nil* or siNT). Cisplatin responses tested in HEI-OC1 were analyzed by 2-way ANOVA at multiple concentrations, or one-way ANOVA at a single concentration of cisplatin, using Bonferroni multiple comparisons test. Time course experiments in HEI-OC1 cells were analyzed by 2-way ANOVA with Dunnett’s multiple comparisons test. Neuromast scores were analyzed via one-way ANOVA with Tukey’s multiple comparisons test. All statistical analyses were performed using Prism 7.2.

## Author Contributions

Designed research: GB AL NMP IKD CD AMR WTA APB

Performed research: GB AL NMP IKD CD AMR

Analyzed data: GB AL NMP IKD CD AMR WTA APB

Wrote the paper, or provide their own description: GB AL NMP AMR WTA APB

## Acknowledgments

This work was supported by operating grants to APB from the Canadian Institutes of Health Research (MY2-155361) and Hair Massacure (FN 2489). This research has also been funded by the Li Ka Shing Institute of Virology (LKSIoV) and the generous support of of the Stollery Children’s Hospital Foundation through the Women and Children’s Health Research Institute (WCHRI). GB was supported by studentships from the University of Alberta Faculty of Medicine & Dentistry and WCHRI, while AL was supported by studentships from the Alberta Cancer Foundation and WCHRI. CD was supported by a studentship from the LKSIoV. APB is a Tier 2 Canada Research Chairholder. AR is an ISAC SRL Emerging Leader. Some experiments were performed at the University of Alberta Faculty of Medicine & Dentistry Flow Cytometry Facility, which receives financial support from the Faculty of Medicine & Dentistry and Canada Foundation for Innovation (CFI) awards to contributing investigators.

## Notes

### Competing Interest Statement

The authors have declared no competing interest.

