## Supplemental Figs 1-3 for "Toll-like receptor 4 is activated by platinum and contributes to cisplatin-induced ototoxicity"

**This PDF file includes:**

Figures S1 to S3

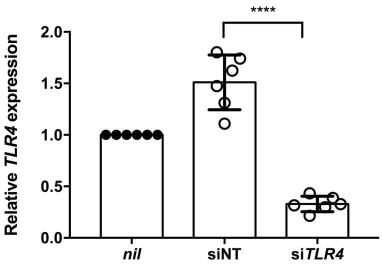


**Fig. S1. *TLR4* expression in HeLa cells is significantly reduced by transient silencing.** Relative *TLR4* expression levels were determined in cisplatin-treated HeLa cells transfected with non-targeting (siNT) or *TLR4*-targeting (si*TLR4*) siRNA molecules. Expression was normalized to untransfected (*nil*) cells with mean and standard deviation are indicated (n=6). Data are from two independent experiments. ****, *P*<.0001 as determined by one-way ANOVA with Dunnet's multiple comparison test.

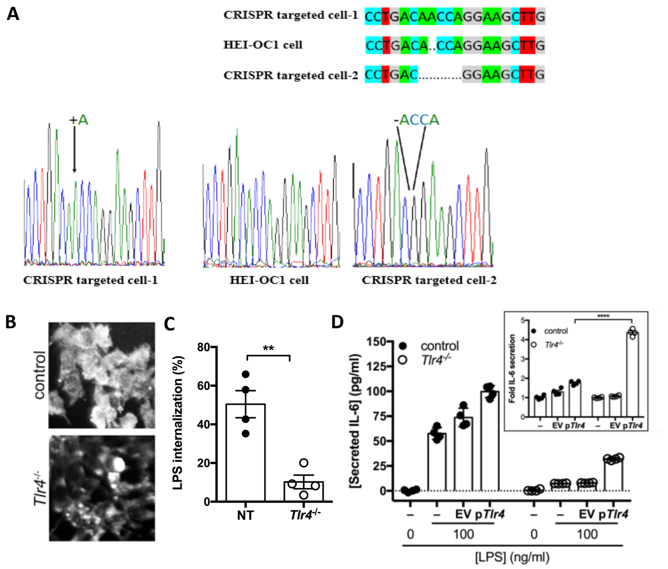

**Fig. S2. *Tlr4*$^{-/-}$ HEI-OC1 cells show diminished LPS-responsiveness unless complemented with *Tlr4*.** Non-targeting control HEI-OC1 cells were compared to Tlr4-targeted CRISPR/Cas9 edited HEI-OC1 cells. **(A)** Genomic DNA from the *Tlr4* locus of *Tlr4-/-* cells was subcloned into plasmids, Sanger sequenced and compared to Sanger sequencing of the *Tlr4* locus in wild type HEI-OC1 cells. Sequences from the *Tlr4*$^{-/-}$ cell line contained a single nucleotide insertion or four nucleotide deletion but no wild type sequence. The results are summarized at the top. **(B)** *Tlr4*$^{-/-}$ and control cells were stained with an anti-TLR4 antibody and visualized by immunofluorescence. **(C)** The internalization of fluorescently conjugated LPS was quantified in *Tlr4*$^{-/-}$ and control cells by flow cytometry. Data (mean ± SD) are from 4 independent experiments. **(D)** *Tlr4*$^{-/-}$ and control cells were transfected with empty vector (EV), *Tlr4* (p*Tlr4*), or left untransfected (-) and the cells were treated with LPS (100 ng/mL). Secreted IL-6 was quantified as a metric for TLR4 activation (n=4). ***Inset,*** fold induction of IL-6 secretion was determined relative to the untransfected cells treated with LPS. Data (mean ± SD) are from 4 independent experiments. **, $P<.01$; ****, $P<.0001$ as determined by unpaired t-test (C) or 2-way ANOVA with Bonferroni multiple comparison test (D).

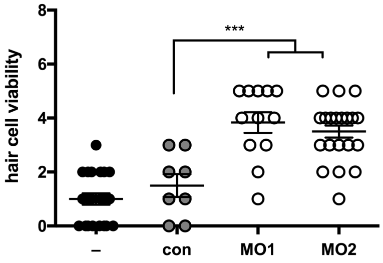


**Fig S3. Cisplatin-induced neuromast toxicity is reduced in independent *tlr4bb* knockdowns.** Fish were pre-treated with control-, and splice-targeting *tlr4bb* (MO1) or translation-targeting *tlr4bb* morpholinos prior to treatment with 15 μM cisplatin. Viable hair cells were enumerated via DASPEI staining and fluorescence microscopy. Each data point (circles) represents a score of hair cell integrity in an individual animal (taken from multiple samples per animal), whereas lines represent mean ± SD (n=8-22). ***, $P<.001$ as determined by ANOVA with Tukey's multiple comparison testing.
